# Scale-invariant geometric data analysis (SIGDA) provides robust, detailed visualizations of human ancestry specific to individuals and populations

**DOI:** 10.1101/431585

**Authors:** Max Robinson, Anat Zimmer, Terry Farrah, Denise E. Mauldin, Nathan D. Price, Leroy E. Hood, Gustavo Glusman

## Abstract

Scale invariance is a common property of physical laws and a key concept in perspective drawing, which aims to provide a meaningful two-dimensional representation of a more complex, three-dimensional scene. Here we describe Scale Invariant Geometric Data Analysis (SIGDA), a new, general exploratory data analysis (EDA) method based on normalization of data to scale invariance. We discuss similarities and differences between SIGDA and two widely-used EDA methods, Correspondence Analysis (CA) and Principal Components Analysis (PCA). We then illustrate SIGDA’s ability to analyze and visualize population structure relationships within the data that inspired its development: genetic marker data, in which context PCA is considered a standard method. We show that SIGDA provides significant advantages over PCA of the same data, including: (a) robust detection and separation of a larger number of population axes, leading to (b) better separation of annotated populations; (c) separation of an independent allele frequency axis interpretable as a proxy for allele age, (d) visualization of marker flow between populations (population *history*), and (d) robust detection and visualization of relationships between closely-related individuals and among family groups. Although this illustration focuses on a specific task, SIGDA is a general-purpose EDA method and derives its advantages from its novel approach to fundamental issues in data analysis, rather than clever sampling or other task-specific methodology.

**One Sentence Summary:** We illustrate the advantages of Scale Invariant Geometric Data Analysis (SIGDA), a new exploratory data analysis method similar to PCA, by applying SIGDA to derive detailed, robust visualizations of the complex history of human population structure from a large sample of single nucleotide variants.

## Overview

Scale-Invariant Geometric Data Analysis (SIGDA) is a new general-purpose exploratory data analysis method with a focus on scale invariance. SIGDA is intended to generalize two widely-used methods which apply to different kinds of data: Principal Components Analysis (PCA) [1,2], which applies z-score normalization to each of a set of random variables (columns) measured on a set of objects (rows), and Correspondence Analysis (CA) [3], which applies a chi-squared model to cross-tabulated counts of observed events (a contingency table). SIGDA interprets each matrix entry as a weight of similarity (or proximity or association) between the containing row and the containing column, or equivalently whatever (hidden) annotation may be associated with each row and column. SIGDA therefore generalizes both PCA and CA by discarding the assumptions which determine their respective approaches to data normalization, and it is SIGDA’s unique approach to data normalization which distinguishes it most from existing methods. SIGDA’s normalization, which we call *projective decomposition*, simultaneously rescales the rows and columns of the data matrix *M* = { *m*_*ij*_ } = { *r*_*i*_*w*_*ij*_*c*_*j*_ } by positive scaling factors *r*_*i*_ for each row and *c*_*j*_ for each column to produce the scale-invariant matrix *W* = {*w*_*ij*_ } (see Methods). These scaling factors cancel to preserve every relative ratio of columns across rows of *M* in *W*, 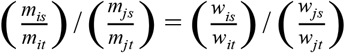, and likewise preserve every relative ratio of rows across columns. SIGDA is therefore designed for data with variation on a scale *relative* to the data’s expected value, while PCA is designed for data varying on an *absolute* scale that is independent of its expected value. Examples of data for which SIGDA’s model may be most appropriate include contingency tables, gene expression data, relative probabilities of events, and other “ratio-scale” data.

Below we illustrate SIGDA’s ability to analyze and visualize complex relationships within the data which inspired its development: patterns of population structure history among the single nucleotide variants observed in the 1000 Genomes Project [4]. PCA is a standard method [5] for visualization of static population structure and encoding population structure numerically for use as a covariate in genome-wide association studies (GWAS). Here we show that SIGDA provides substantially more robust and detailed analyses of population structure from the same data. We also demonstrate SIGDA’s ability to visualize relationships between the scale invariant dimensions and an explicit scale dimension. In genetic marker data, scale is proportional to minor allele frequency, and is a proxy for time as the expected age of the derived marker allele [6]. We therefore argue that SIGDA visualizations of genetic marker data describe human population divergence and admixture over time.

## Materials and Methods

### Overview

SIGDA’s goal is to represent both the rows and the columns of an *m*×*n* data matrix as points in a shared coordinate system; the process by which this is accomplished (Figure 1) is described in the three phases of simultaneous capture of two views of the data with by projective decomposition (Capture), ordination of axes of variation within the captured views (Orient), and construction of a merged image in a single, shared coordinate space (Project).

**Figure 1.**
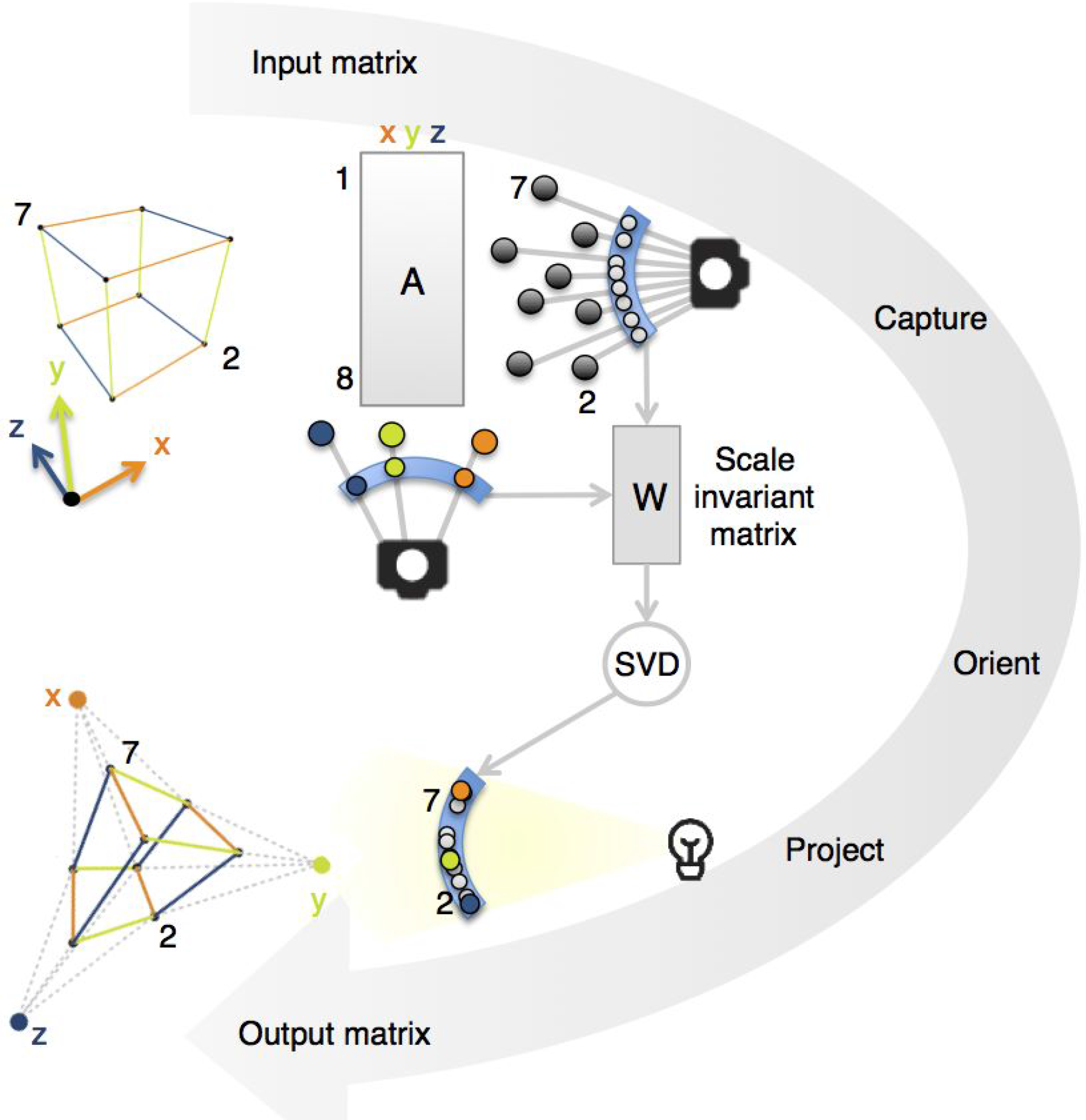
Overview of SIGDA. *Input matrix:* The corners of a cube (points, upper left; numbered) with edges (lines) parallel to the coordinate axes x, y, and z (arrows) are defined in an 8 × 3 data matrix A (Supplementary Table S1). *Capture:* SIGDA interprets A twice: as 3D points defined by the eight rows (dark gray, right of A), and unconventionally (colored, below A) as an 8-dimensional point for each axis (column). Conceptually, projective decomposition simultaneously “focuses” these row and column points onto spheres (blue arcs); procedurally, it rescales each row and column of A to form a scale-free matrix W. *Orient:* SVD identifies axes of variation as min{8,3} = 3 pairs of orthogonal vectors, the orientation of each axis as it appears in the two views. Using these corresponding axes, the two sets of points superimpose onto a single spherical surface in a shared coordinate system (bottom center). *Project:* SIGDA flattens the spherical merged image by projection onto a flat hyperplane, removing one dimension. *Output matrix:* The result is an 11 × 2 matrix (11=8+3, 2 = min {8,3}−1). These points form a 2D drawing (lower left) of the cube plus the 3 column points. Extended edges (dashed) intersect at the column points to representing the parallel coordinate axis, confirming that SIGDA has created a 3-point perspective diagram, with the column points representing points at infinity on each original coordinate axis.

### Normalization by projective decomposition

SIGDA therefore interprets each matrix entry twice: as a coordinate in a row vector, and as a coordinate in a column vector (Fig. 1). Acting as a kind of stereoscopic camera viewing the rows and columns from the origin, projective decomposition (as summarized above) iteratively rescales the row vectors to have length 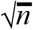 and the column vectors to have length 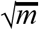, “focusing” the vectors onto two focal spheres. The rows of the normalized matrix *W* identify *m* “row points” forming an image of the data matrix on the sphere of radius 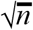 in *R*^*n*^, while the columns of *W* identify *n* “column points” forming an alternative image of the same data on the sphere of radius 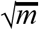 in *R*^*m*^. Each of these images consists of a set of points on a hypersphere embedded in a higher-dimensional Euclidean space; the points do not vary by distance from the origin, only on their direction from the origin. Furthermore, both images are embedded in subspaces of at most *k* = *min* {*m*, *n*} dimensions (either more than *k* points are embedded in *R*^*k*^, or *k* points are embedded in a higher-dimensional space, and therefore contained in the subspace spanned by the *k* vectors from the origin and to the *k* points themselves).

### Ordination of axes

SIGDA determines the “relative orientation” between these two *k*-dimensional subspaces by singular value decomposition (SVD) [7], obtaining *k* pairs of corresponding singular vectors. Each singular vector pair represents the same axis of variation among the data in *W* as it appears in the row image and the column image, allowing the origins and singular vectors of the two subspaces to be equated and interpreted as a shared coordinate space (*R*^*k*^). The row and column points therefore occupy the spheres of radius 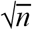 and 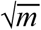, respectively, in *R*^*k*^.

### Projection to a shared coordinate system

SIGDA projects the spherical row and column images onto the flat hyperplane a distance 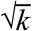 from the origin along the principal axis of variation (corresponding to the first singular vector pair), with three effects: (1) elimination of the dimension along the principal axis of variation, and rescaling the row points from a curved surface with radius of curvature 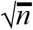 and the column points from a curved surface with radius of curvature 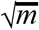 to the same hyperplane, with the effect of (2) eliminating the difference in scale between the row points and the column points, and (3) placing all points into a geometrically flat space, one with a Euclidean geometry. The projected row and column therefore merge into a shared, (*k* − 1)-dimensional Euclidean coordinate system, one in which geometrical patterns have their familiar, Euclidean characteristics, simplifying their interpretation. The order of the axes of this space is inherited from the singular value decomposition of *W*, and therefore in order of decreasing singular value (as in PCA).

### SIGDA’s output

We refer to these scale-invariant axes as *EC*_1_, *EC*_2_, …, *EC*_*k*−1_ (EC stands for “Euclidean Coordinate” (or “Espalier Coordinate”, see below). ECs are not simple linear combinations of the columns of *M*, and are therefore not “components” as defined by PCA despite the apparent similarities. Each EC axis is scale invariant; for analysis of relationships between these (*k* − 1) scale invariant dimensions and an appropriate notion of scale, SIGDA uses the mean absolute value of each row (column) in *M* as the appropriate scale dimension.

### Mathematical and implementation details for projective decomposition

Projective decomposition separates the *m*×*n* data matrix *A* = { *a*_*ij*_ } = *f*_*A*_*D*_α_*WD*_β_ into a product of scaling factors (*f*_*A*_, the Frobenius norm of *A*, and the positive row factors α and column factors β on the diagonals of the diagonal scaling matrices *D*_α_ and *D*_β_, respectively) and the normalized matrix 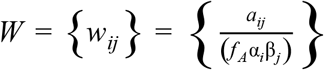, where *W* and each of its rows and columns are constrained to have unit root-mean-square (RMS). Projective decomposition can be performed efficiently by standard matrix balancing algorithms [8,9] adapted to balance RMS instead of row and column sums. *W* is invariant^1^ to transposition and scaling of *A*, and we therefore refer to *W* as the *scale invariant* matrix similar^2^ to *A*.

Unlike PCA’s scale standardization, projective decomposition is not translation invariant: adding a value to any row or column of the data matrix in general alters the direction from the “camera” to some or all of the data points, yielding a different normalized matrix. Whenever the origin lies inside the convex hull of the row or column data points, the image SIGDA produces will conceptually surround a point at infinity (the observer) with the data, and appear “inside out”. Unless this is a desired effect, a constant may be added to the matrix to ensure the origin lies outside the data. Scale standardization also standardizes the observer’s view of the data, which is often considered an advantage; however the ability to control the observer’s view can resolve whether geometric properties are intrinsic to the data, i.e. independent of the chosen view.

### SIGDA Visualizations

Scale invariance is a central concept of projective geometry, the mathematics of perspective drawing. We illustrate that SIGDA produces perspective visualizations with the analysis of an 8×3 matrix representing the corners of a cube (Fig. 1, lower left; Supplementary Table S1). The merged image is a perspective visualization of the cube containing a point for each row and a point for each column; the column points, each of which represents a coordinate axis in the data matrix, become vanishing points (points at infinity along the coordinate axes) at which all lines parallel to the coordinate axes intersect. However, in general SIGDA will be used on data with many more than 3 dimensions, and this interpretation as a perspective drawing is therefore of limited utility. This connection with projective geometry is, however, at the heart of our “data camera” analogy.

Analysis of a matrix whose rows contain weighted sums of three vectors *a*, *b*, and *c* (Figure 2, Supplementary Table S2) illustrate some of SIGDA’s scale invariance properties. Note that the points for rows containing weighted sums (computed as labeled) lie on line segments connecting the points for the pair of vectors they combine in SIGDA’s results (Fig. 2b), but not PCA’s (Fig. 2a). Geometrically, a line segment represents every weighted *average* of its endpoints, and a weighted average is a scale invariant representation of a class of weighted sums. Looking again at the PCA analysis, the points for weighted sums can be seen as displaced from collinearity in proportion total weight. These observations are consistent with complete separation of a scale dimension from *k* − 1 scale invariant EC axes by SIGDA, but retention of at least some scale effects in the *k* PC axes of PCA.

**Figure 2.**
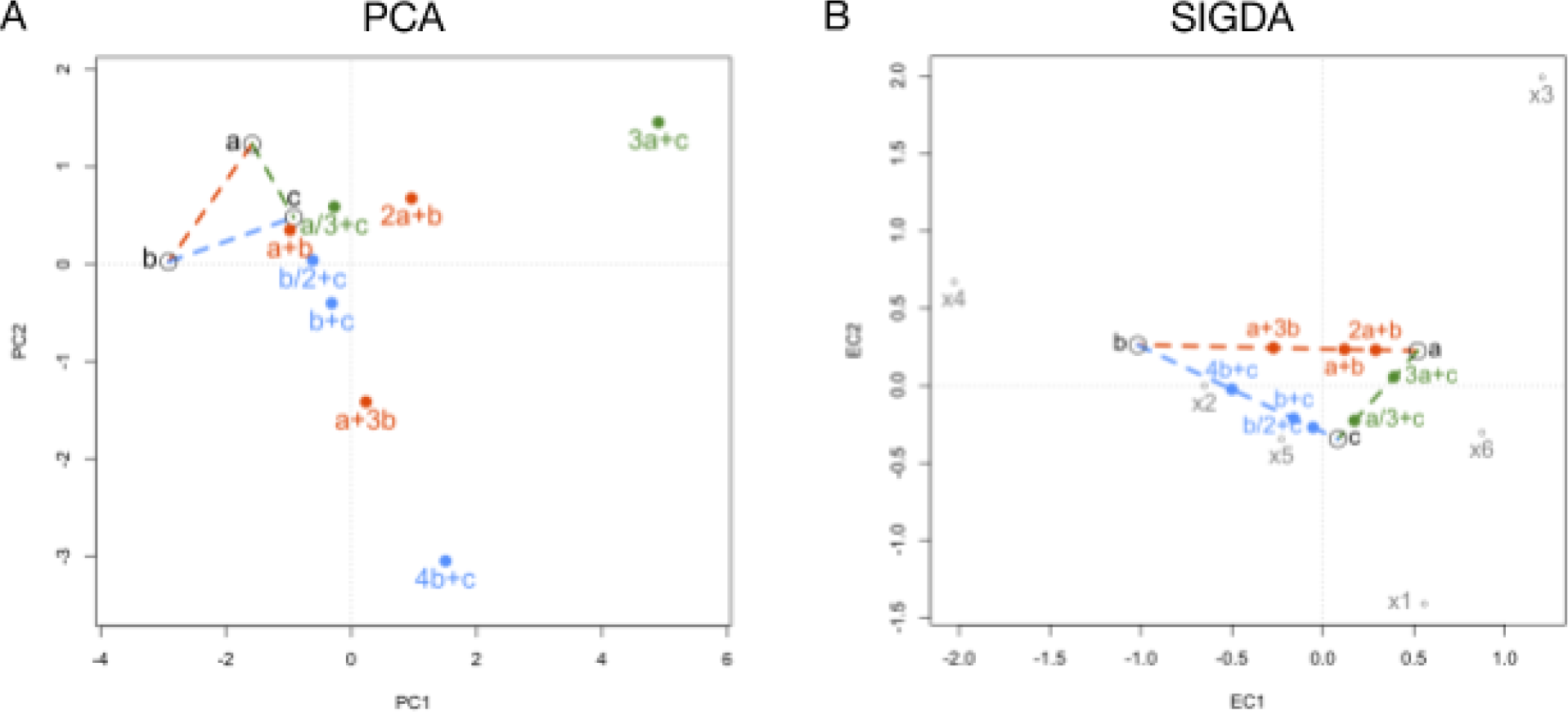
SIGDA represents scale invariant relationships geometrically. We analyzed an 11 × 6 matrix (Supplementary Table S2) containing 3 independent vectors (a, b, and c, open circles) and weighted sums (filled circles) of a and b (3 sums, dashed line; red), a and c (2 sums, dashed line; green), or b and c (3 sums, dashed line; blue). (A) Biplot of the first two principal components from a PCA analysis. PCA places the weighted sums at a distance from the associated vectors proportional to total weight, preserving scale relationships between rows despite standardizing column scales; roughly collinear points are not all linearly related. (B) Biplot of the first two dimensions (ECs) from the a SIGDA analysis. SIGDA places the point for each weighted sum directly between the points for the associated vectors, preserving the scale invariant linear relationships among all sets of sums. The rows for vectors a, b, and c were randomly generated, and no association between the row points and column points (x1-x6, gray) is expected or apparent.

SIGDA’s behavior on a data matrix representing a four-dimensional nonlinear relationship, the Ideal Gas Law *PV* = *R nT* (Fig. 3), illustrates a valuable property of scale invariant analysis. To construct this dataset (Supplementary Table S3), we randomly generated 2000 combinations of pressure *P*, volume *V*, and mass *n*, and computed temperature according to the Ideal Gas Law, 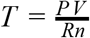. Stereograms of PCA results (Fig. 3a) and SIGDA results (Fig. 3b) are qualitatively different. While the PCA analysis reflects uniform sampling of a rectangular volume of pressure, volume and mass values, the SIGDA analysis is interpretable as a 3D perspective visualization: a saddle surface in three scale-invariant dimensions, bounded by points at infinity along the *P*, *V*, *n*, and *T* axes. The saddle shape is indicative of the algebraic form of the Ideal Gas Law: orthogonal (additive) quadratic terms *P* × *V* and *n* × *T* with opposite signs, α *PV* − β *nT* = 0, where the formal constants α, β represent scale invariance.

**Figure 3.**
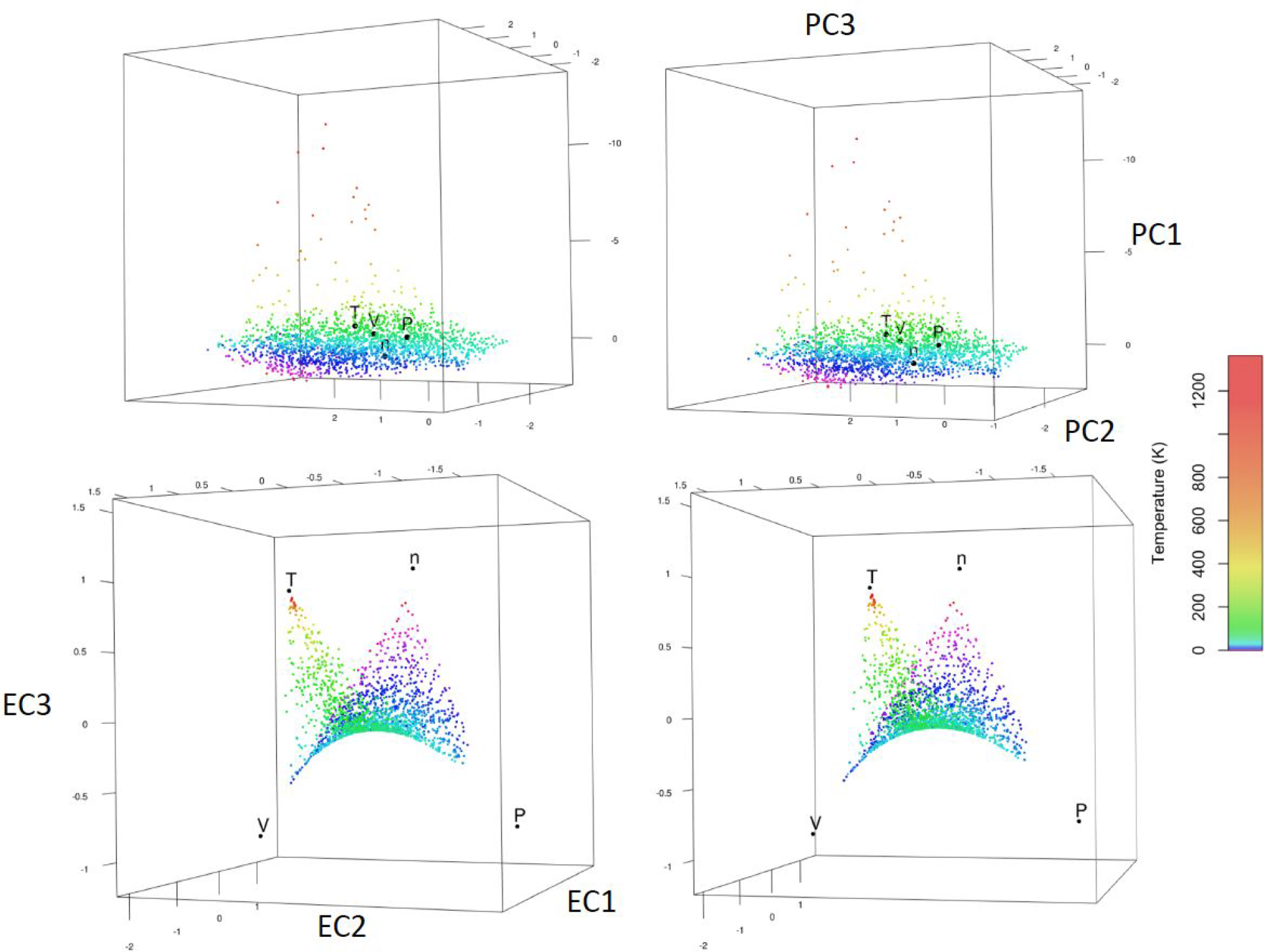
SIGDA reveals the scale invariant geometry of nonlinear relationships. We generated 2000 random sets of pressure (*P*), volume (*V*), and mass (*n*) values and computed the corresponding temperature (*T*) from the Ideal Gas Law, *T* = *PV* /*Rn* (SI units; *R* = 0.082057 L atm mol-1 K^−1^) to produce a 2000×4 matrix (Supplementary Table S3). Stereo image pairs of the first three dimensions produced by PCA (A) and SIGDA (B) contain a point for each row, colored by temperature on a logarithmic scale. The SIGDA images also contain a point representing each variable (B, black points). The PCA triplot (A) shows that temperature is associated with PC3. SIGDA (B) produces a smooth saddle surface (*z* = *x*^2^ − *y*^2^), with the point for each variable a point at infinity corresponding to high values of that variable (compare colors, location of *T*). As annotated, this saddle directly represents the algebraic structure of the Ideal Gas Law, *EC*3 = *PV* − *nT*.

### Espalier plots

SIGDA records an explicit scale dimension, the mean value within each row and column. In prior work [10], we defined an Espalier plot to have a scale independent horizontal axis and scale as the vertical axis; the name refers to garden espaliers (trellises) used to simplify the complex, inaccessible branching structure of fruit trees and other plants readily accessible on a two-dimensional, vertical surface.

### Genetic marker data

We conceived of SIGDA during analysis of single nucleotide variant (SNV) data from the 1000 Genomes Project’s phase 3 cohort [4]. This data set comprises whole genome sequences, assembled, corrected, and filtered by rigorous comparison of multiple data sources and diverse experimental methods, for 2504 DNA samples from 26 annotated populations, including over 20 million biallelic autosomal SNVs with 5 or more observations of the minor allele (MAF >= 5/5008 = 0.1%). We randomly labeled these SNVs with a number from 1 to 50, creating 50 partitions of approximately equal size (414,652 to 417,011 SNVs), and formed a sparse genotype matrix for each subset. Each matrix entry is the count (0, 1, or 2) of occurrences of the minor allele for a SNV observed in each DNA sample; in subset 01, 11.7% of the genotype matrix entries were nonzero. We focused on analysis of subset 01; the remaining subsets were reserved for robustness analysis (below).

The typical preparation of genetic markers for PCA analysis of population structure retains only single nucleotide polymorphisms (SNPs, SNVs with MAF > 5%) that are considered “ancestry-informative” by an information-theoretic criterion [11]. However, since a genotype matrix contains counts of observed minor alleles, MAF corresponds to SIGDA’s scale axis: two alleles are counted per sample, and the average value in a column is therefore twice the MAF of the corresponding SNV. We therefore excluded only SNVs with MAF too low to accurately estimate (fewer than 5 minor alleles observed). Our focus in this paper is not on efficient use of marker data; it is on SIGDA as a general exploratory data analysis tool.

## Results

We evaluated SIGDA’s utility for population structure analysis by comparing PCA and SIGDA analyses of a large set of genetic variants: 415724 SNVs data from the 1000 Genomes Project’s phase 3 cohort (subset 01; see Methods). The PC_2_ - PC_1_ biplot (PCA, Fig. 4A) and the corresponding EC_2_ - EC_1_ biplot (SIGDA, Fig. 4B) of the 2504 samples (dots), colored by annotated population (see legend) are broadly similar: both organize most samples near the corners or edges of an AFR-EAS-EUR triangle typically visible in PCA visualizations of worldwide population structure. The samples farther into the triangle’s interior on both biplots are generally from populations acknowledged to be admixed (the AMR populations and AFR populations ASW and ACB), but note that the SAS populations are also within the triangle, and are simply separated from the other continental populations on axes with lower singular values.

**Figure 4.**
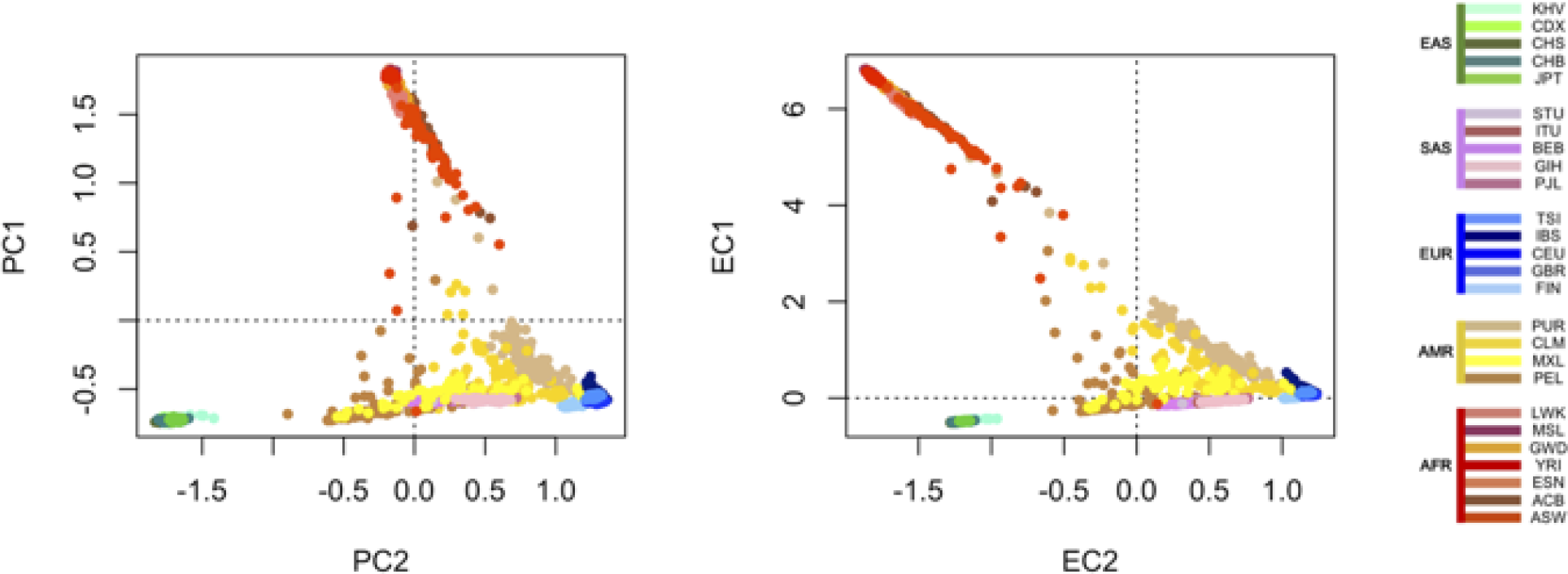
Organization of rows (samples). Both PCA and SIGDA produce coordinates for rows (samples) as well as columns (SNVs). In PCA, row coordinates are interpreted as the location of a geometric point for each sample, but column coordinates are interpreted as the direction of a line through the origin (axis) for each SNV; in SIGDA both row coordinates and column coordinates are interpreted as geometric points. The plots above show a point for each sample at its corresponding row coordinates; the column coordinates, which would be shown as vectors in a traditional PCA plot, are too numerous and are omitted for clarity. These plots are therefore a traditional PCA biplot (left) of the 2,504 samples of the 1000 Genomes cohort shows one point for each row (sample), and the corresponding SIGDA biplot (right). Each sample is colored by population (legend), and spatial organization of samples reflects population structure. Population structure is broadly similar between the two approaches on these first two axes, with subtle differences. SIGDA, but not PCA, separates AFR, EUR, SAS, and EAS into different quadrants. SIGDA also produces sharper linear structures.

Two notable differences between these PC biplots and EC biplots with points representing the samples are (a) the position of the origin and (b) the relative location of EAS and AFR samples on PC_2_ (AFR < EAS < EUR) versus EC_2_ (EAS < AFR < EUR). Together, these differences cause AFR, EUR, SAS, and EAS samples to segregate to different quadrants of the SIGDA plot, while AFR is located on the PC_1_ axis and quadrant 4 of Figure 4A contains both EUR and SAS. The four continental populations are therefore partitioned by EC thresholds at 0 in the EC biplot, but not by PC thresholds at 0. In subsequent dimensions (not shown), the order and degree of separations among the 26 annotated populations remain similar through the first 5 to perhaps as many as 10 dimensions (depending on the standards applied), then increasingly diverge [^3^ A more thorough analysis of these patterns has been performed and will be described in a future version of this manuscript. For now, we only claim that the nature of the differences in subsequent dimensions are similar to those observable in the biplots we have shown.]. Thus, SIGDA provides roughly the same analysis of population structure provided by PCA, with some differences in the order of separations between closely related populations in the context of a diverse, worldwide cohort.

SIGDA also provides EC coordinates and a scale coordinate for each column (variant; see Figure 5 A,B,D). In this SNV data, and under the simplifying assumption that the more frequent (major) allele is ancestral while the less frequent is derived, SIGDA’s scale values are directly proportional to variant minor allele frequency (MAF). As discussed above (see Methods), MAF is a proxy for allele age, and to the extent that EC coordinates reflect elements of population structure (the previous paragraph demonstrated that they do in a manner similar to PC coordinates), an Espalier plot therefore depicts an aspect of population structure specific to the chosen EC plotted against a proxy for evolutionary time. We therefore examined the meaning of a population structure Espalier plot in more detail.

**Figure 5.**
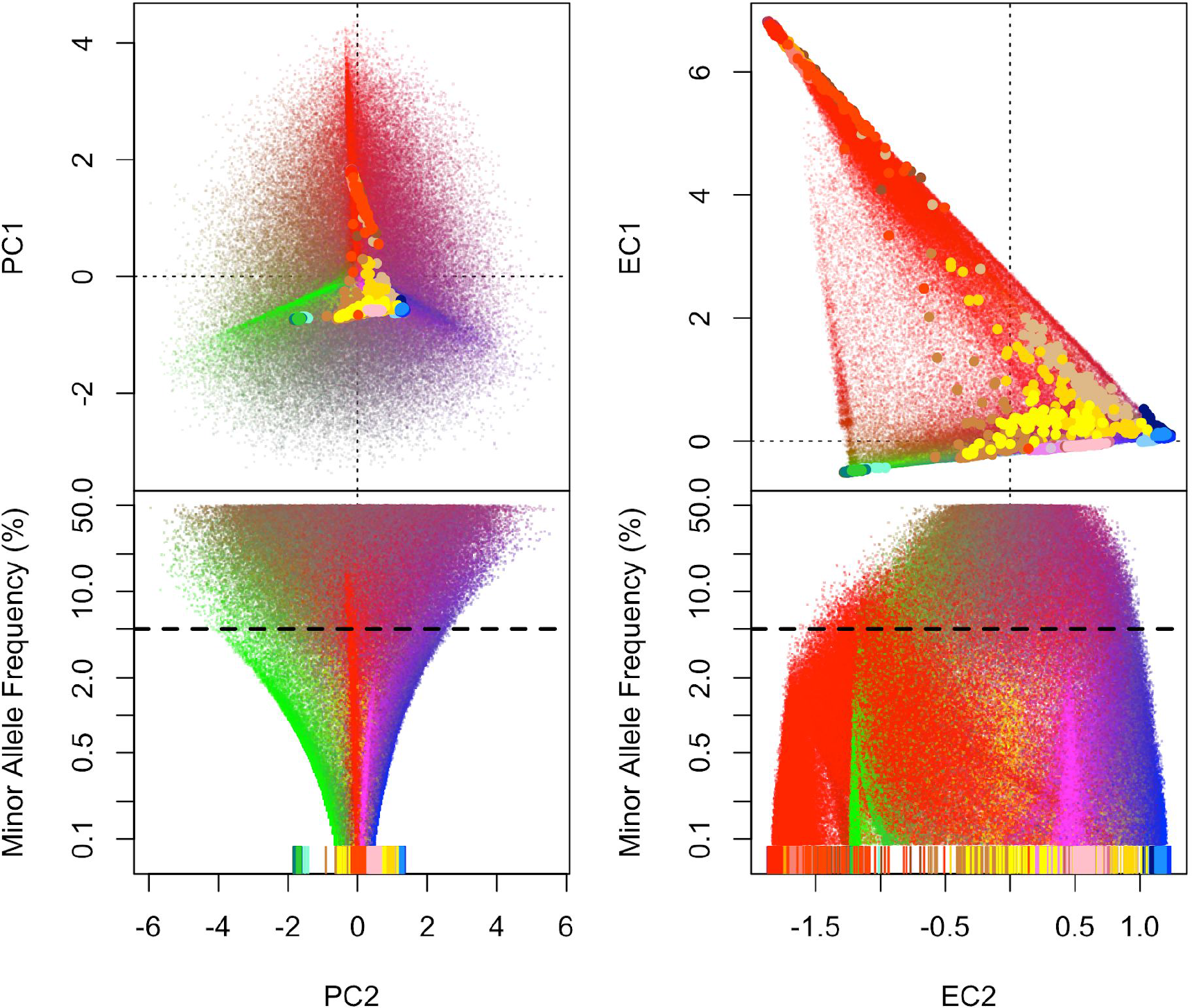
Organization of columns (SNVs). Both PCA and SIGDA produce coordinates for columns (SNVs) as well as rows (samples). In a SIGDA biplot (upper right), both rows (large points; located and colored as in Figure 4) and columns (small points) are represented as points in the same coordinate system (Espalier Coordinates, ECs). Column points have been added to the PCA biplot (upper left) for comparison; in PCA, these values are interpreted as loadings (weights) that relate the new component axes (PCs) to the original column axes (SNVs). Thus, in PCA the column coordinates correspond to geometric vectors rather than points as they are shown here. The color of each column point representing a variant (all four plots) is determined by the distribution of the minor allele of the corresponding SNV among the five continental areas (AFR, AMR, EAS, EUR, and SAS; see Methods). Column point colors are computed so that a SNV whose minor alleles are only observed in AFR samples is pure red (R=1, G=0, B=0); only observed in EAS is pure green (R=0, G=1, B=0); only observed in EUR, pure blue (R=0, G=0, B=1); only observed in AMR, yellow (R=1, G=1, B=0); only observed in SAS, purple (R=1, G=0, B=1). The lower plots show the organization of SNVs by PCA (lower left) and SIGDA (lower right) on PC_2_ (x-axis, shared with upper plot) by minor allele frequency (y-axis, logarithmic scaling). Rare minor alleles are low on the y-axis, common minor alleles are above. The dashed line indicates a minor allele frequency of 5%; SNVs with a frequency below 5% are often excluded from population structure analyses by PCA. Only the EC or PC coordinate of the rows are indicated in the lower plots (vertical lines near x-axes; compare to dots of same color in upper plots).

It is not possible to make a direct comparison between Espalier plots produced by these two methods: PCA neither singles out scale as a separate dimension, nor are principal components scale-invariant dimensions. Espalier plots are specific to SIGDA’s analysis methodology. We nevertheless compared plots of variant loadings against MAF (Figure 5C) as the most direct analogue of SIGDA’s Espalier plots provided by PCA.

In the Espalier and Espalier-like plots (Fig. 5D, 5C) the position of each sample along the x-axis is shown (colored vertical lines near the x-axis) in the same population-specific colors shown in the color bar accompanying Figure 4. The points representing SNVs, which do not belong to annotated populations, are assigned roughly corresponding colors according to the frequency with which the minor allele was observed among individuals from the five continental areas (AFR, AMR, EAS, EUR, and SAS) as an <R, G, B> color value determined as follows: R is the fraction of the SNV’s minor alleles observed among samples from AFR, AMR, or SAS; G is the fraction among samples from AMR or EAS; and B is the fraction among samples from EUR or SAS. This has the effect that pure red (<1, 0, 0>) corresponds to AFR samples alone, and likewise yellow (<1, 1, 0>) represents AMR, green EAS, blue EUR, and purple SAS, with SNVs observed on multiple continents assigned intermediate colors.

These two color schemes expose scale-invariant (vertical) patterns in the SIGDA Espalier plot (Fig. 5D): samples from each continent are vertically aligned with monochromatic “cones” of rare variants whose minor alleles are only observed among samples from the aligned continent, confirming that the SNVs are organized horizontally by population information. It is well-accepted that MAF provides a rough estimate of allele age (how long a derived allele has been segregating in a population [6], with rare variants (more precisely, rare derived alleles) expected to have arisen more recently than common variants. Thus, the vertical dimension in the Espalier plots shown here, MAF, can be considered a proxy for time. We defer further interpretation of the vertical and diagonal patterns apparent in Fig. 5D to the Discussion section.

The EC values SIGDA provides for each individual variant support an analysis similar to a previously described “chromosome painting” strategy [12], but with MAF as an additional dimension. As a proof of principle, we therefore identified the minor alleles present on Chromosome 2 of selected individuals, and visualized them in the context of the full variant sample (Figure 6). In accordance with our observations in Figure 5, we painted the chromosome with only rare variants (MAF < 5%; lower 3 figures), with the corresponding Espalier plots for reference (upper figures). We found that different ECs provided separation of rare variants apparently inherited from distinct genetic backgrounds by an admixed individual (NA19657 from population MXL), whose genome contains sets of rare variants shared predominantly among distinct genetic backgrounds (red, African ancestors; yellow, likely Native American ancestors; blue, European ancestors). We note that the same kind of analysis is not well-supported by PC coordinates produced by PCA, as variant loadings are confounded mixtures of scale and population-related information. We note that the sample of SNVs analyzed here are only 2% of those available, and even restricting variants to MAF < 5%, this analysis could be done at 50x the resolution along the chromosome if all available variants were used.

**Figure 6.**
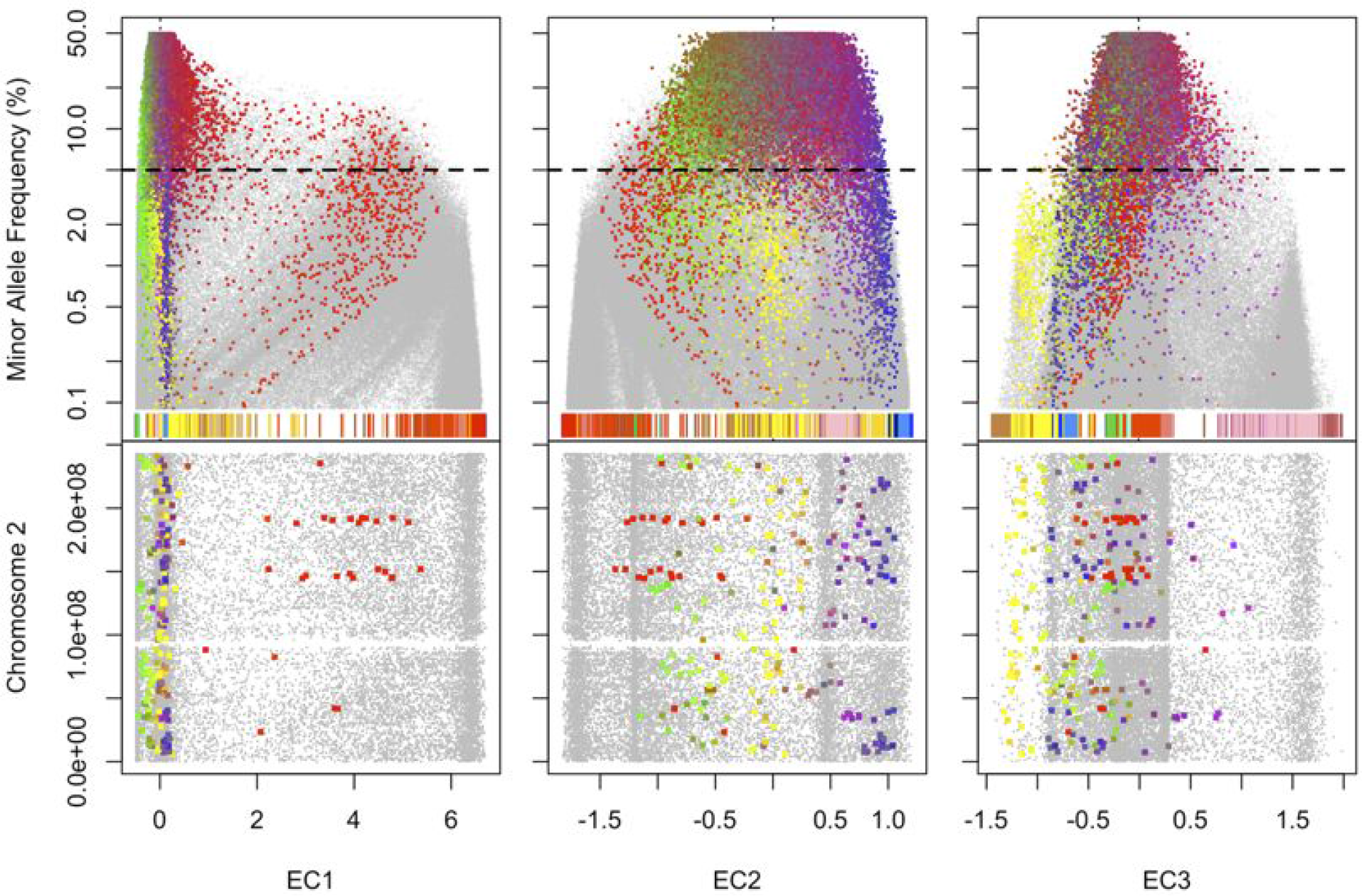
“Chromosome painting” visualization of admixture in a single individual along Chromosome 2. Variants for which MXL sample NA19657 bears the minor allele (colored points) are shown on SIGDA’s first three principal axes (EC1, EC2, EC3), providing a map of the differing origin of chromosomal segments in a recently admixed individual. All analyzed variants are shown for context (gray points). The color of each variant encodes the relative association of the minor allele with individuals from the different continental areas (see Methods), minor alleles found only in AFR individuals are red; exclusively AMR alleles are yellow; EAS, green; EUR, blue; SAS, purple. These colors are broadly consistent with thematic organization of colors assigned to individuals in each population (colored vertical bars near the x-axis, upper plots). *Upper plots:* all analyzed variants are shown by minor allele frequency (MAF, y-axis). *Lower plots:* recent variants (MAF <5%) on Chr. 2 are shown by position along the chromosome (y-axis). Each EC shown correlates with a historical separation of populations: EC1 (left) correlates with the out-of-Africa event (out-of-AFR < AFR); among the non-AFR populations, EC2 (center) correlates with separation of the indo-European populations (EUR, SAS) from EAS; EC3 (right) correlates with separation within the indo-European populations (AMR+EUR vs. SAS). In the lower plots, horizontal position correlates with alleles common in AFR, EUR, and AMR (colors) in contiguous segments. Diagonal patterns (upper plots) represent variants present in a chromosomal segment that is common in a (source) population, but rare in a (destination) population. In such a segment, common variants (high MAF) are present in many individuals in the source population, but few in the destination population; rare variants (low MAF) were present in a smaller fraction of the source population but in the same number of recipients in the destination population as common variants in the same chromosomal segment, and are therefore more equally represented in the source versus destination populations, leading to a gradual shift in horizontal position correlated to MAF.

In addition to the direct application of providing information about the location and size of chromosomal segments on Chromosome 2 this individual likely inherited from different ancestral populations, Figure 6 clearly demonstrates the time-dependent, multi-dimensional nature of population structure, as well as the value of accounting for the age of the derived allele, even through a proxy such as minor allele frequency, in the assessment of population structure via genetic variation.

## Discussion

We have introduced SIGDA, a general-purpose, scale-invariant approach to exploratory data analysis of numerical matrix data more similar to PCA, CA, and other geometric ordination techniques [13–16] than to other dimensionality reduction techniques [17–21]. This method applies directly to non-negative data matrices, including but not limited to contingency tables, genetic marker data, measurements of absolute physical quantities such as pressure, temperature, volume, mass, and a wide variety of other scientific measurements. We illustrated SIGDA’s behavior and usefulness on genetic marker data, an excellent example of high-dimensional data set contain a rich set overlapping and hierarchical patterns with direct interpretations. We have applied SIGDA to a variety of other measured datasets, and have observed that the advantages of SIGDA highlighted here are relevant in many different settings.

The goal of exploratory data analysis is to separate and expose for analysis the patterns contained in a set of data early in the analysis of the data, when those patterns may not yet be known. While PCA, CA, and SIGDA use similar mathematical tools for this common purpose, we believe a brief comparison of the goals of each method will clarify the settings in which SIGDA can be most effective. Briefly, PCA was originally introduced (by Pearson [1] and Hotelling [2]) as a method for aligning coordinate axes along mutually independent linear trends in the data. Geometrically, PCA treats the rows of a data matrix as points, and both the columns and the linear trends (principal axes or components) as lines. In contrast, CA [3] is frequently used in ecology to compare two categorizations, one into rows and one into columns, of a set of counted observations (a contingency table). CA represents each row and each column as a geometric point in an abstract space; points representing corresponding categories have corresponding coordinates, while contrasting categories are separated to distant regions of space.

SIGDA synthesizes ideas present in CA and PCA, bringing the geometric correspondence strategy of CA to the broader class of datasets typically analyzed with PCA. SIGDA achieves this synthesis through its focus on scale and scale invariance, which is embodied in normalization by projective decomposition. SIGDA also takes advantage of connections between scale invariance and projective geometry to project the correspondences among and between rows and columns into a more familiar Euclidean space, greatly increasing the relevance of our geometric intuition and providing an interpretation of the resulting visualizations as high-dimensional perspective drawings.

Each of the examples we have provided illustrate novel, unique analytical capabilities provided by SIGDA: multipoint perspective visualization of high-dimensional data; visualization of geometric, rather than statistical relationships; enhanced interpretation of row-row, row-column, and column-column relationships; and a familiar geometric framework to guide interpretations. In population structure analysis, where PCA is already a standard method, we have highlighted two particular advantages: (1) by placing the column points in the same coordinate system as the row points, SIGDA organizes the SNVs by population in a manner directly comparable to the organization of samples, allowing the population information carried by SNVs to be interpreted in the same way population information for samples is currently understood; and (2) improving the interpretability of both scale-independent population information (populations which endure over time) and the scale dependence of population information (changes in population structure over time) by separating scale from scale-independent axes.

It is important to note that both of these advantages generalize to other settings: placing rows and columns in the same coordinate system enriches our ability to interpret both irrespective of the field from which the data is drawn, and it is difficult to imagine a dataset in which our notion of scale, the average values in each row and column, will fail to represent some directly interpretable aspect of the phenomenon under study. We therefore anticipate that SIGDA will be a useful, general purpose approach to exploratory data analysis.

## Acknowledgments

We would like to thank Nigel Clegg, who helped implement and test software related to this work. SIGDA is available at https://github.com/PriceLab/SIGDA. This work was supported by the Inova Translational Medicine Institute, NIH grants P50 GM076547 and U54 EB020406, and the MJ Murdock Charitable Trust.

## Supplementary Materials

**Table S1:** Coordinates of a cube with edges aligned to the coordinate axes.

|  | "x" | "y" | "z" |
| --- | --- | --- | --- |
| "1" | 4 | 3 | 2 |
| "2" | 14 | 3 | 2 |
| "3" | 4 | 13 | 2 |
| "4" | 14 | 13 | 2 |
| "5" | 4 | 3 | 12 |
| "6" | 14 | 3 | 12 |
| "7" | 4 | 13 | 12 |
| "8" | 14 | 13 | 12 |

**Table S2:** Linear combinations of three 6D vectors.

|  | "x1" | "x2" | "x3" | "x4" | "x5" | "x6" |
| --- | --- | --- | --- | --- | --- | --- |
| "a" | 0.413 | 0.501 | 0.570 | 0.041 | 0.470 | 0.934 |
| "b" | 0.009 | 0.447 | 0.029 | 0.489 | 0.237 | 0.007 |
| "c" | 0.819 | 0.904 | 0.141 | 0.325 | 0.811 | 0.996 |
| "a+3b" | 0.440 | 1.842 | 0.657 | 1.509 | 1.180 | 0.955 |
| "2a+b" | 0.834 | 1.449 | 1.169 | 0.572 | 1.177 | 1.875 |
| "a+b" | 0.422 | 0.948 | 0.599 | 0.531 | 0.707 | 0.941 |
| "4b+c" | 0.856 | 2.692 | 0.257 | 2.281 | 1.758 | 1.025 |
| "b+c" | 0.828 | 1.351 | 0.170 | 0.814 | 1.048 | 1.003 |
| "b/2+c" | 0.824 | 1.128 | 0.156 | 0.569 | 0.930 | 1.000 |
| "3a+c" | 2.057 | 2.408 | 1.851 | 0.449 | 2.222 | 3.797 |
| "a/3+c" | 0.956 | 1.071 | 0.331 | 0.338 | 0.968 | 1.307 |

**Table S3: Simulated data conforming to the Ideal Gas Law.**

See PVnRT.txt at https://github.com/PriceLab/SIGDA.

## Footnotes

1 Scaling: If *A* = *f*_*A*_ *D*_α_ *W D*_β_ and *D*_α_, *D*_λ_ are *m×m* matrices with positive diagonals and *D*_α_, *D*_λ_ are *n×n* matrices with positive diagonals, then *V* = *D*_λ_ *A D*_ϱ_ = *f*_*V*_ *D*_λα_ *W D*βρ, where the diagonals *λ α* 0 and *β*_ϱ_ > 0 are positive. Transposition: 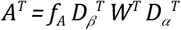.

2 Two *m×n* matrices *A* and *B* are said to be *similar* if strictly positive scaling vectors *γ* of length *m* and *ϱ* of length *n* exist that satisfy the equation *B* = *D*_γ_ *A D*_ϱ_, where as above *D*_*x*_ = *Diag*{*x*}, the square matrix with diagonal *x*.

## References

1. Pearson K. L III. On lines and planes of closest fit to systems of points in space. The London, Edinburgh, and Dublin Philosophical Magazine and Journal of Science. Taylor & Francis; 1901;2: 559–572.

2. Hotelling H. Analysis of a complex of statistical variables into principal components. J Educ Psychol. Warwick & York; 1933;24: 417.

3. Greenacre M. Correspondence Analysis in Practice, Second Edition. 2007.

4. 1000 Genomes Project Consortium, Auton A, Brooks LD, Durbin RM, Garrison EP, Kang HM, et al. A global reference for human genetic variation. Nature. 2015;526: 68–74.

5. Price AL, Patterson NJ, Plenge RM, Weinblatt ME, Shadick NA, Reich D. Principal components analysis corrects for stratification in genome-wide association studies. Nat Genet. 2006;38: 904–909.

6. Kimura M, Ohta T. The age of a neutral mutant persisting in a finite population. Genetics. 1973;75: 199–212.

7. Golub G, Kahan W. Calculating the Singular Values and Pseudo-Inverse of a Matrix. Journal of the Society for Industrial and Applied Mathematics Series B Numerical Analysis. Society for Industrial and Applied Mathematics; 1965;2: 205–224.

8. Sinkhorn R, Knopp P. Concerning nonnegative matrices and doubly stochastic matrices. Pacific J Math. Mathematical Sciences Publishers; 1967;21: 343–348.

9. Knight P, Ruiz D, Uçar B. A Symmetry Preserving Algorithm for Matrix Scaling. SIAM J Matrix Anal Appl. Society for Industrial and Applied Mathematics; 2014;35: 931–955.

10. Robinson M, Eley G, Vockley JG, Niederhuber JE and Glusman G. Espaliers: A Visualization Method for Big Data. JSM Proceedings, Statistical Computing Section. Alexandria, VA: American Statistical Association; 2015. pp. 2897–2096.

11. Rosenberg NA, Li LM, Ward R, Pritchard JK. Informativeness of genetic markers for inference of ancestry. Am J Hum Genet. 2003;73: 1402–1422.

12. Lawson DJ, Hellenthal G, Myers S, Falush D. Inference of population structure using dense haplotype data. Copenhaver GP, editor. PLoS Genet. Public Library of Science; 2012;8: e1002453.

13. Kruskal JB. Toward a practical method which helps uncover the structure of a set of multivariate observations by finding the linear transformation which optimizes a new “index of condensation.” Statistical Computation. 1969. pp. 427–440.

14. Hochreiter S-A. Clevert D, Obermayer K. A new summarization method for affymetrix probe level data. Bioinformatics. 2006;22: 943–949.

15. Wang C, Mahadevan S. Manifold alignment using Procrustes analysis. Proceedings of the 25th international conference on Machine learning - ICML ’08. 2008. doi:10.1145/1390156.1390297

16. Alter O, Brown PO, Botstein D. Singular value decomposition for genome-wide expression data processing and modeling. Proc Natl Acad Sci U S A. 2000;97: 10101–10106.

17. Sammon JW. A Nonlinear Mapping for Data Structure Analysis. IEEE Trans Comput. 1969;C-18: 401–409.

18. Tenenbaum JB, de Silva V, Langford JC. A global geometric framework for nonlinear dimensionality reduction. Science. 2000;290: 2319–2323.

19. Roweis ST. Nonlinear Dimensionality Reduction by Locally Linear Embedding. Science. 2000;290: 2323–2326.

20. Coifman RR, Lafon S, Lee AB, Maggioni M, Nadler B, Warner F, et al. Geometric diffusions as a tool for harmonic analysis and structure definition of data: diffusion maps. Proc Natl Acad Sci U S A. 2005;102: 7426–7431.

21. Platzer A. Visualization of SNPs with t-SNE. PLoS One. 2013;8: e56883.

